# ChAdOx1 nCoV-19 (AZD1222) protects Syrian hamsters against SARS-CoV-2 B.1.351 and B.1.1.7

**DOI:** 10.1101/2021.03.11.435000

**Authors:** Robert J. Fischer, Neeltje van Doremalen, Danielle R. Adney, Claude Kwe Yinda, Julia R. Port, Myndi G. Holbrook, Jonathan E. Schulz, Brandi N. Williamson, Tina Thomas, Kent Barbian, Sarah L. Anzick, Stacy Ricklefs, Brian J. Smith, Dan Long, Craig Martens, Greg Saturday, Emmie de Wit, Sarah C. Gilbert, Teresa Lambe, Vincent J. Munster

## Abstract

We investigated ChAdOx1 nCoV-19 (AZD1222) vaccine efficacy against SARS-CoV-2 variants of concern (VOCs) B.1.1.7 and B.1.351 in Syrian hamsters. We previously showed protection against SARS-CoV-2 disease and pneumonia in hamsters vaccinated with a single dose of ChAdOx1 nCoV-19. Here, we observed a 9.5-fold reduction of virus neutralizing antibody titer in vaccinated hamster sera against B.1.351 compared to B.1.1.7. Vaccinated hamsters challenged with B.1.1.7 or B.1.351 did not lose weight compared to control animals. In contrast to control animals, the lungs of vaccinated animals did not show any gross lesions. Minimal to no viral subgenomic RNA (sgRNA) and no infectious virus was detected in lungs of vaccinated animals. Histopathological evaluation showed extensive pulmonary pathology caused by B.1.1.7 or B.1.351 replication in the control animals, but none in the vaccinated animals. These data demonstrate the effectiveness of the ChAdOx1 nCoV-19 vaccine against clinical disease caused by B.1.1.7 or B.1.351 VOCs.

## Main

The COVID-19 pandemic produced an unprecedented development of SARS-CoV-2 vaccines, and just over a year after the beginning of the outbreak a total of 12 vaccines have been authorized or approved globally. As the pandemic progressed, several variants of concern (VOCs) have been detected. These include the B.1.1.7 and B.1.351 VOCs. The B.1.1.7 VOC was first detected in the United Kingdom and has seven amino acid (AA) substitutions and two deletions in the spike protein^1,2^ compared to the original Wuhan isolate, Wuhan-Hu-1. The B.1.351 VOC was first detected in South Africa and has eight AA substitutions and one deletion in the spike protein^3^ (Table 1). All currently licensed vaccines are based on the spike protein of Wuhan-Hu-1, thus, concerns have been raised that the presence of these changes may affect vaccine efficacy. The goal of this study was to evaluate ChAdOx1 nCoV-19 (AZD1222) vaccine efficacy in Syrian hamsters, when challenged using naturally occurring isolates of the VOCs B.1.1.7 and B.1.351.

**Table 1.**
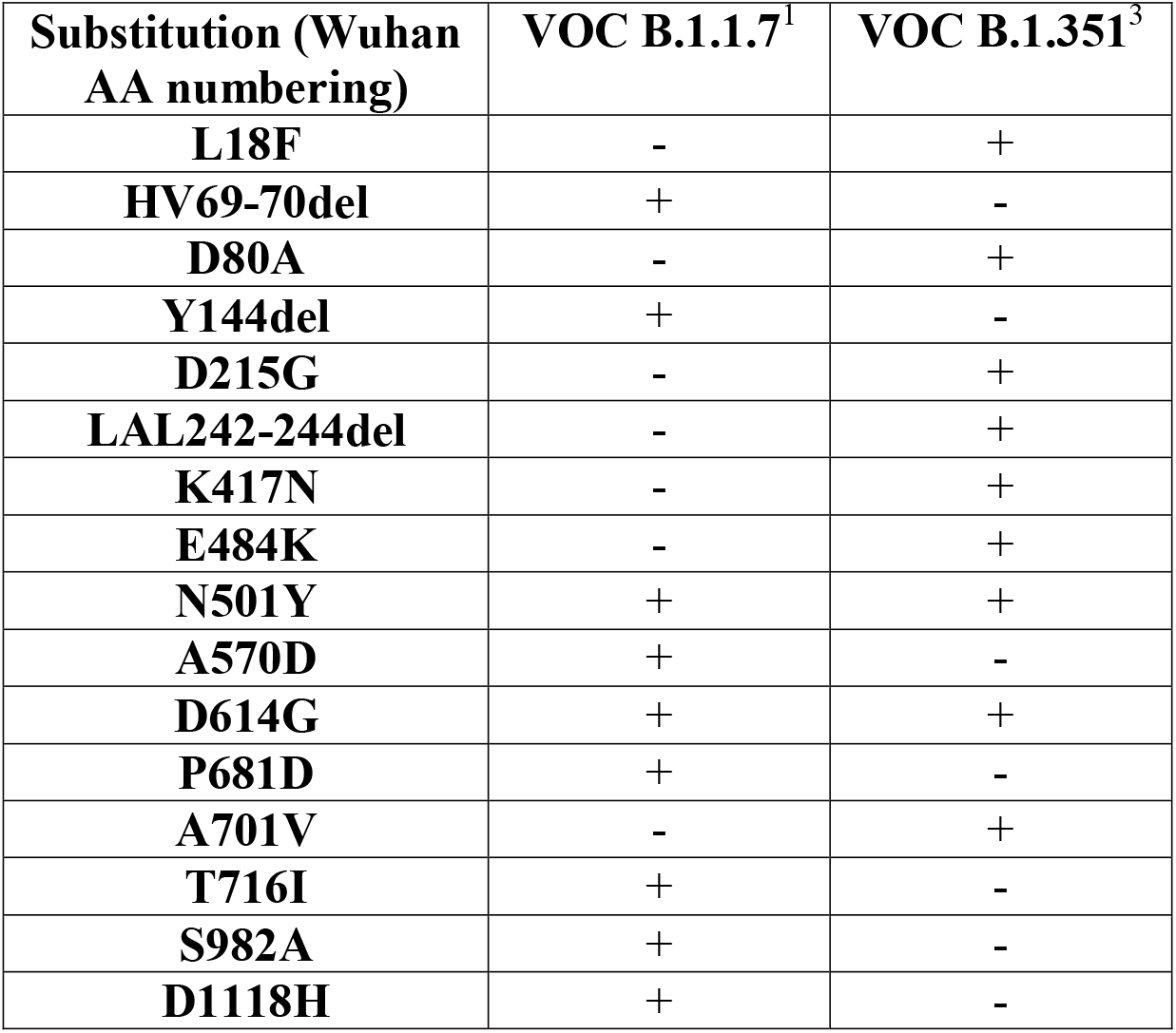
AA substitutions detected in the spike protein of VOCs B.1.1.7 (EPI_ISL_601443) and B.1.351 (EPI_ISL_678615) compared to Wuhan-Hu-1 (NC_045512).

Syrian hamsters (N=10 per group) were vaccinated intramuscularly with either ChAdOx1 nCoV-19 or ChAdOx1 green fluorescent protein (GFP, 2.5 × 10^8^ IU/hamster) 30 days prior to intranasal challenge with SARS-CoV-2. Vaccination with ChAdOx1 nCoV-19 resulted in high titers of binding antibodies against the SARS-CoV-2 full-length spike protein and receptor binding domain (Figure 1a) at 25 days post vaccination. We then investigated neutralizing antibody titers in serum against infectious virus. Neutralization of B.1.351 was significantly reduced compared to neutralization of B.1.1.7 (Figure 1b, mean titer of 15 vs 142, p < 0.0001, Mann-Whitney test).

**Figure 1.**
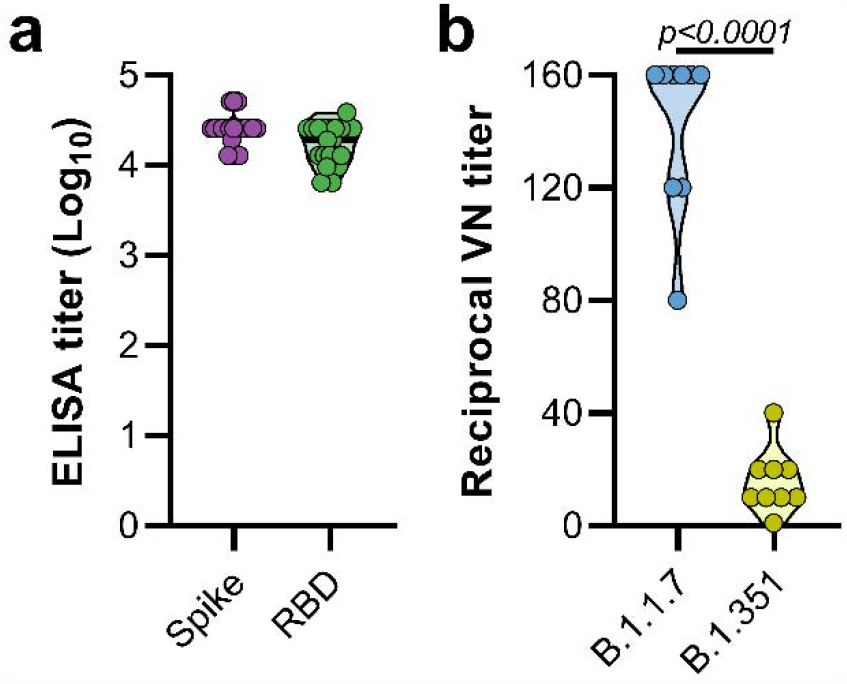
Vaccination of Syrian hamsters with ChAdOx1 nCoV-19 elicits binding and neutralizing antibodies against B.1.1.7 and B.1.351. a. Violin plot of binding antibodies against spike protein or RBD of SARS-CoV-2 (clade A) in serum obtained 25 days post vaccination with ChAdOx1 nCoV-19. b. Violin plot of virus neutralizing antibody titers against B.1.1.7 or B.1.351 in serum obtained 25 days post vaccination with ChAdOx1 nCoV-19. Statistical significance determined via Kruskall-Wallis test.

### ChAdOx1 nCoV-19 vaccinated hamsters are protected against lower respiratory tract infection with B.1.1.7

Hamsters were inoculated with B.1.1.7 via the intranasal route. Weight loss was observed in control hamsters whereas vaccinated hamsters continued to gain weight throughout the experiment (Figure 2a). A significant difference in weight between vaccinated and control hamsters was observed starting at 4 days post infection (DPI) for B.1.1.7 (Figure 2a, Student’s t-test corrected for multiple comparisons using the Holm-Šidák method) and continued throughout the remainder of the experiment. Four out of ten hamsters per group were euthanized at 5 DPI and lung tissue was harvested. Lung:body weight ratios on 5 DPI were significantly lower in vaccinated animals compared to control animals (Figure 2b, p=0.0286, Mann-Whitney test), indicating no or reduced pulmonary edema in ChAdOx1 nCoV19-vaccinated animals. Lung tissue of all control animals contained high levels of sgRNA (Figure 2c, 10^10^ copies/gram tissue), and was comparable to sgRNA levels previously detected in lung tissue of control animals challenged with SARS-CoV-2 D614G (hCoV-19/USA/MT-RML-7/2020)^4^. Conversely, no sgRNA was detected in lung tissue obtained from vaccinated hamsters challenged with B.1.1.7 (Figure 2c, p=0.0286, Mann-Whitney test). High levels of infectious virus were detected in lung tissue of all control animals, whereas no vaccinated animals had detectable infectious virus in lung tissue (Figure 2d, p=0.0286, Mann-Whitney test).

**Figure 2.**
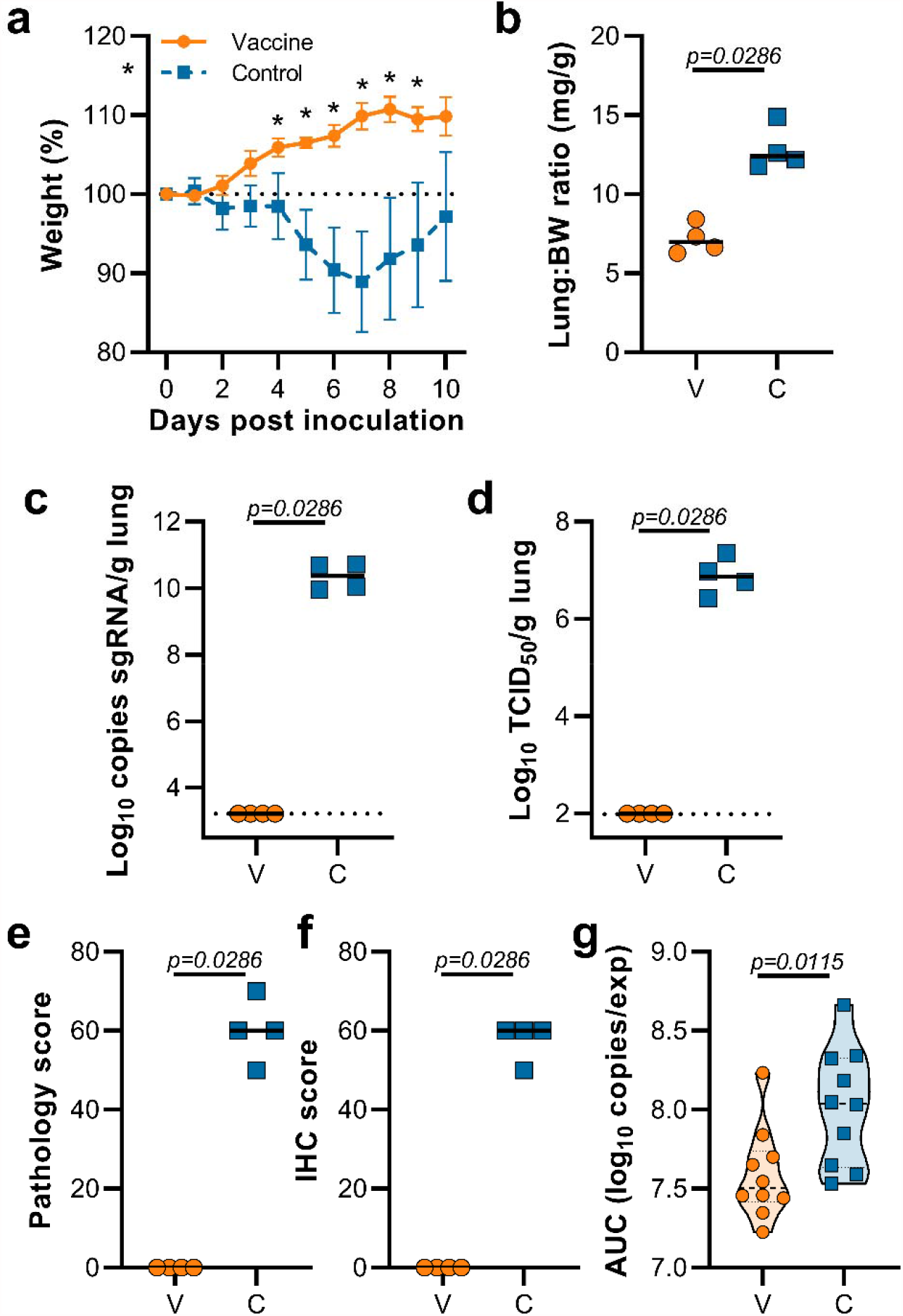
Vaccination of Syrian hamsters with ChAdOx1 nCoV-19 prevents lower respiratory tract infection with SARS-CoV-2 VOC B.1.1.7. a. Relative weight upon intranasal challenge with 10^4^ TCID_50_ of B.1.1.7. Shown is geometric mean with 95% confidence interval (CI). * = p-value<0.05, corrected for multiple comparisons using the Holm-Šidák correction. b. Lung:body weight (BW) ratio (mg:g) of hamsters euthanized at 5 DPI. Line = median.c. sgRNA viral load in lung tissue obtained at 5 DPI. Line = median. Dotted line = limit of detection. d. Infectious SARS-CoV-2 titer in lung tissue obtained at 5 DPI. Line = median. Dotted line = limit of detection. e. Percentage affected lung tissue per animal as determined via histology. Line = median. f. Percentage of lung tissue positive for SARS-CoV-2 antigen per animal. Line = median. g. Truncated violin plot of area under the curve (AUC) analysis of shedding as measured by sgRNA analysis in swabs collected on 1 – 5 dpi. Dashed line = median. Dotted line = quartiles. Statistical significance determined via mixed-effect analyses (a), or Mann-Whitney test (b-g). V = ChAdOx1 nCoV-19 vaccinated; C = ChAdOx1 GFP vaccinated; Orange circle = Hamsters vaccinated with ChAdOx1 nCoV-19, Blue square = Hamsters vaccinated with ChAdOx1 GFP.

Lung tissue was then evaluated for histology. The percentage of lung tissue that showed pathology and the percentage of lung tissue that was positive for SARS-CoV-2 antigen was determined by a veterinary pathologist blinded to the study group allocations. Whereas no pathology nor SARS-CoV-2 antigen was found in lung tissue of vaccinated animals, this was abundantly present in lung tissue of control animals (Figure 2e,f). Finally, oropharyngeal swabs were collected on 1 to 5 DPI, evaluated for sgRNA, and an area under the curve was calculated per animal to determine the total amount of virus shed. We observed a significant decrease in the total amount of virus found in oropharyngeal swabs from vaccinated animals compared to control animals (Figure 2g, Mann-Whitney test).

Lung tissue was then evaluated for histology. Microscopically, pulmonary lesions of control animals consisted of a moderate to marked broncho-interstitial pneumonia extending into the adjacent alveoli previously observed in hamsters inoculated with SARS-CoV-2 WA1 or a D614G isolate^4,5^. Bronchi and bronchioles had multifocal necrotic epithelial cells and moderate numbers of infiltrating neutrophils and macrophages. Alveolar septa were expanded by edema fluid and leucocytes. In contrast, vaccinated animals did not show any evidence of SARS-CoV-2 pathology (Figure 3a-d). Immunohistochemistry using a monoclonal antibody against SARS-CoV-2 demonstrated viral antigen in bronchial and bronchiolar epithelium, type I and II pneumocytes as well as pulmonary macrophages within the control animals, but not in vaccinated animals (Figure 3e-f).

**Figure 3.**
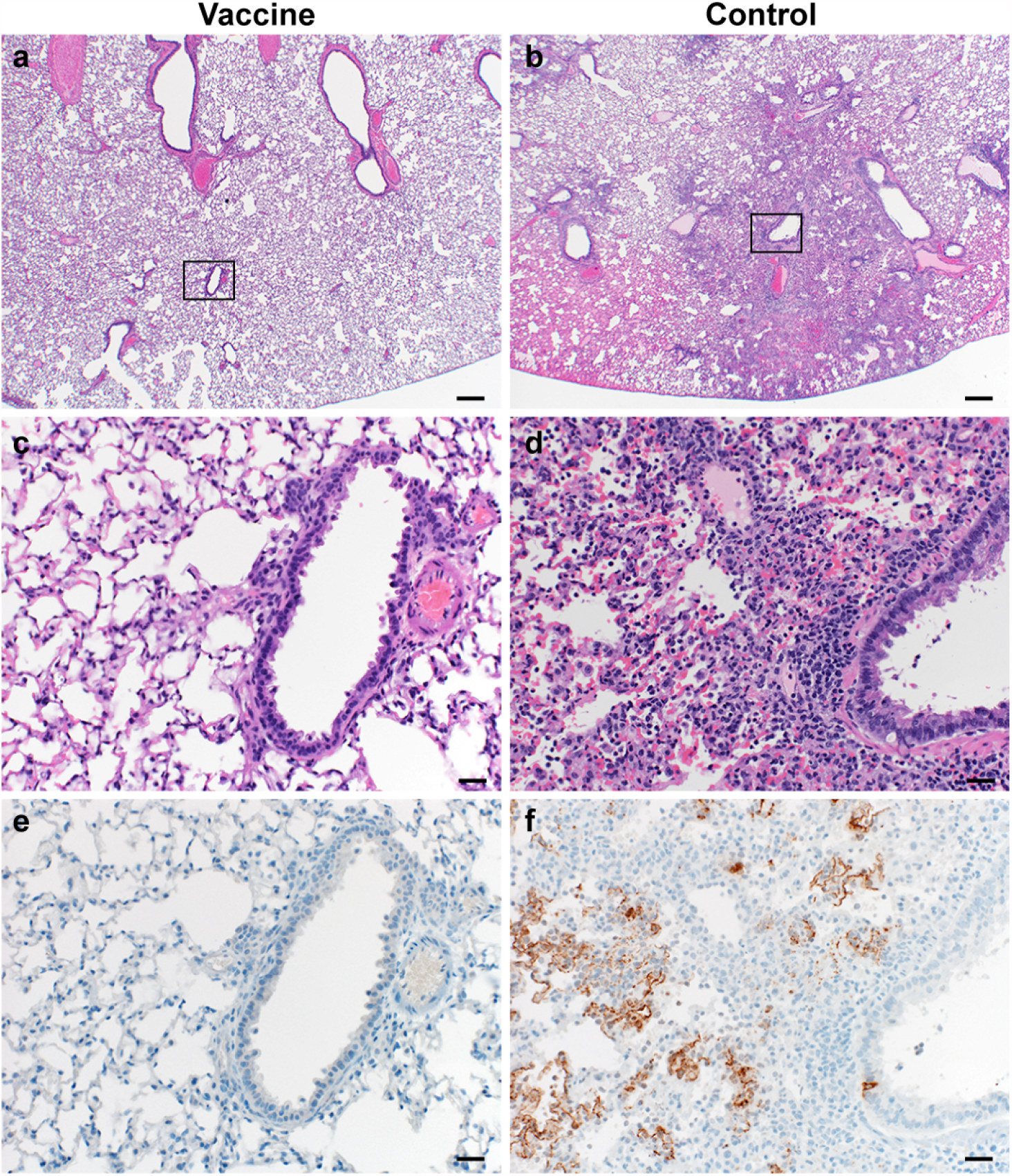
Pulmonary effects of direct intranasal challenge with SARS-CoV-2 variant B.1.1.7 in Syrian hamsters at 5 DPI. a-b. H&E staining, 20x; a. No pathology. b. Focally extensive areas of bronchointerstitial pneumonia. c-d. H&E staining, 200x; c. No pathology. d. Bronchointerstitial pneumonia with alveolar histiocytosis, fibrin and edema. e-f. IHC staining against N protein SARS-CoV-2 (brown). e. No staining. f. Staining of bronchiolar epithelial cells, type I&II pneumocytes and rare macrophages.

### ChAdOx1 nCoV-19 vaccinated hamsters are protected against lower respiratory tract infection with B.1.351

This experiment was repeated using B.1.351 (isolate hCoV-19/South Africa/KRISP-K005325/2020) instead of B.1.1.7 as an inoculation virus. Two AA substitutions were found in the spike protein of the B.1.351 virus stock; Q677H (present at 88%) and R682W (present at 89%). A lack of weight gain was observed in control hamsters, but not vaccinated hamsters, which was significant starting at 6 DPI (Figure 4a, Student’s t-test corrected for multiple comparisons using the Holm-Šidák method). Four out of ten hamsters per group were euthanized at 5 DPI and lung tissue was harvested. Lung:body weight ratios were significantly lower in vaccinated animals compared to control animals (Figure 4b, p=0.0286, Mann-Whitney test).

**Figure 4.**
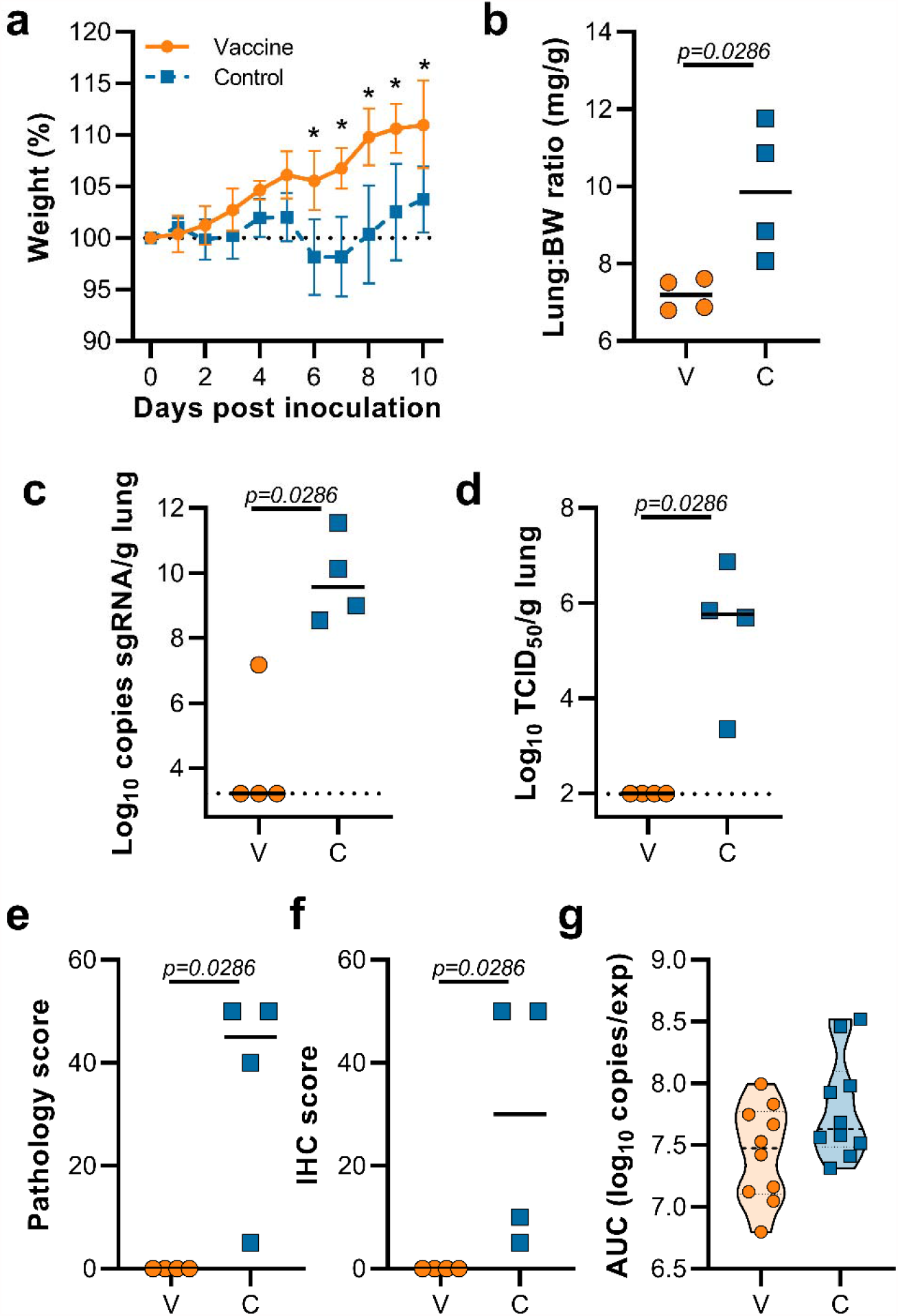
Vaccination of Syrian hamsters with ChAdOx1 nCoV-19 prevents lower respiratory tract infection with SARS-CoV-2 VOC B.1.351. a. Relative weight upon intranasal challenge with 10^4^ TCID_50_ of B.1.351. Shown is geometric mean with 95% confidence interval (CI). * = p-value<0.005, corrected for multiple comparisons using the Holm-Šidák correction. b. Lung:body weight (BW) ratio (mg:g) of hamsters euthanized at 5 DPI. Line = median.c. sgRNA viral load in lung tissue obtained at 5 DPI. Line = median. Dotted line = limit of detection. d. Infectious SARS-CoV-2 titer in lung tissue obtained at 5 DPI. Line = median. Dotted line = limit of detection. e. Percentage affected lung tissue per animal as determined via histology. Line = median. f. Percentage of lung tissue positive for SARS-CoV-2 antigen per animal. Line = median. g. Truncated violin plot of area under the curve (AUC) analysis of shedding as measured by sgRNA analysis in swabs collected on 1 – 5 dpi. Dashed line = median. Dotted line = quartiles. Statistical significance determined via mixed-effect analyses (a), or Mann-Whitney test (b-g). V = ChAdOx1 nCoV-19 vaccinated; C = ChAdOx1 GFP vaccinated; Orange circle = Hamsters vaccinated with ChAdOx1 nCoV-19, Blue square = Hamsters vaccinated with ChAdOx1 GFP.

Lung tissue of all control animals contained high levels of sgRNA, but only one vaccinated animal had relative low levels of sgRNA in lungs (Figure 4c, p=0.0286, Mann-Whitney test). Likewise, high levels of infectious virus were detected in lungs of control animals, but not in lungs of vaccinated animals (Figure 4d, p=0.0286, Mann-Whitney test). As in the previous experiment, no pathology nor SARS-CoV-2 antigen was found in lung tissue of vaccinated animals compared to control animals (Figure 4e,f, p=0.0286, Mann-Whitney test). No decrease in the total amount of virus found in oropharyngeal swabs from vaccinated animals compared to control animals was found (Figure 4g, Mann-Whitney test).

We investigated the presence of the AA substitutions Q677H and R682W observed in 88-89% of the spike protein of our B.1.351 stock in swabs and lung tissue obtained from control animals challenged. Whereas we did find these substitutions in swabs obtained at 1 DPI, they were not present in swabs obtained at 5 DPI. Likewise, the substitutions were only found in lung tissue of one out of four control hamsters (Table 2).

**Table 2.** Presence of substitutions Q677H and R682W in swabs and lung tissue of hamsters directly inoculated with B.1.351.

| AA substitutions | Presence in swabs<br>(1 DPI, N=5) | Presence in swabs<br>(5 DPI, N=3) | Presence in lungs<br>(N=4) |
| --- | --- | --- | --- |
| Q677H | 44.1-65.7% | 0% | 0 (N=3), 81% (N=1) |
| R682W | 44.9-65.8% | 0% | 0 (N=3), 81% (N=1) |

Lung tissue was then evaluated for histology. Microscopically, pulmonary lesions of control animals were comparable to results obtained from previous SARS-CoV-2 infections of hamsters. In contrast, vaccinated animals did not show any evidence of SARS-CoV-2 pathology (Figure 5a-d). Likewise, immunohistochemistry demonstrated viral antigen present in bronchial and bronchiolar epithelium, type I and II pneumocytes as well as pulmonary macrophages within the control animals, but not in vaccinated animals (Figure 5e-f).

**Figure 5.**
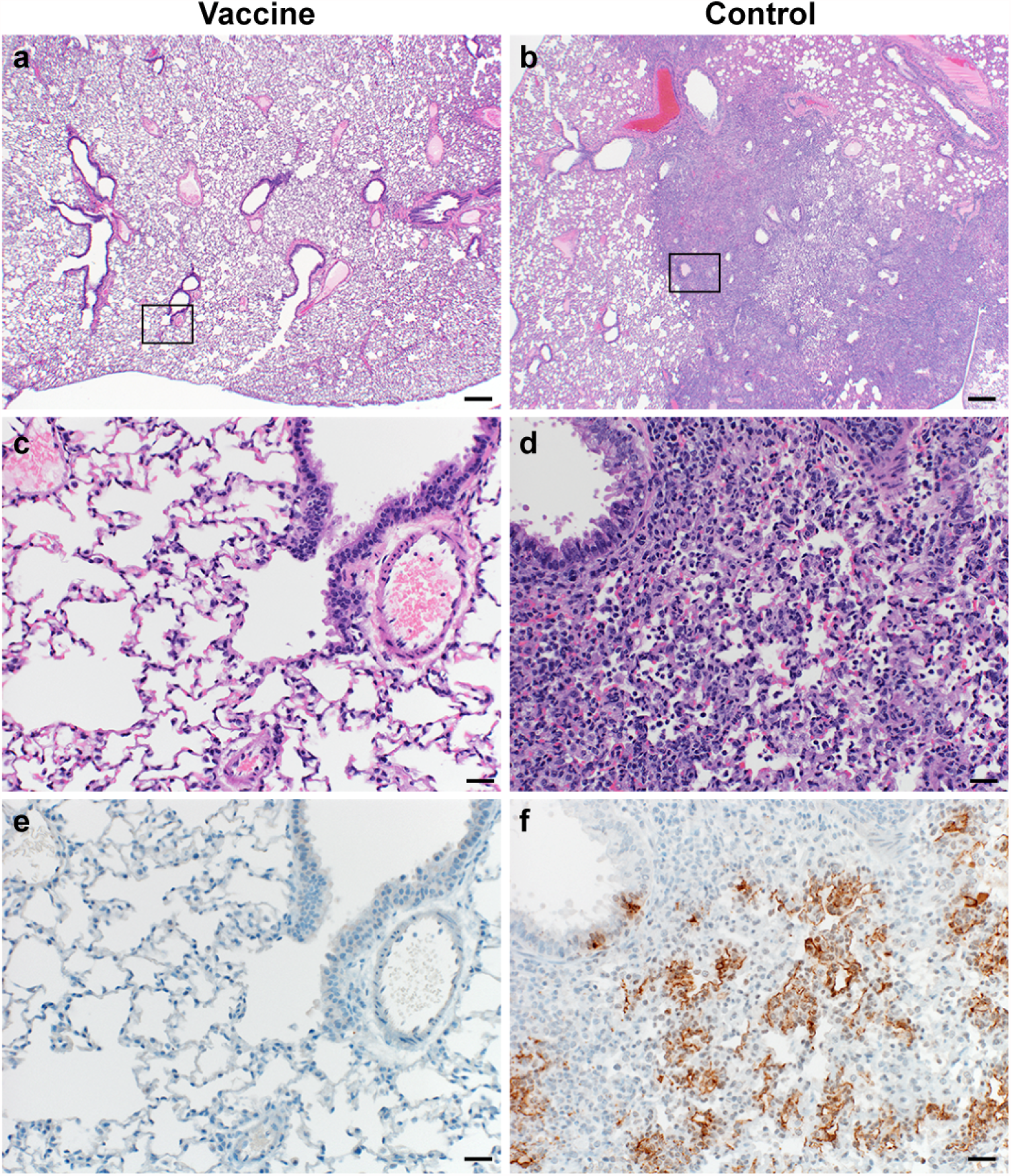
Pulmonary effects of direct intranasal challenge with SARS-CoV-2 variant B.1.351 in Syrian hamsters at 5 DPI. a-b. H&E staining, 20x; a. No pathology. b. Focally extensive areas of bronchointerstitial pneumonia. c-d. H&E staining, 200x; c. No pathology. d. Bronchointerstitial pneumonia with alveolar histiocytosis, fibrin and edema. e-f. IHC staining against SARS-CoV-2 (brown). e. No staining. f. Staining of bronchiolar epithelial cells, type I&II pneumocytes and rare macrophages.

### Hamsters vaccinated with ChAdOx1 nCoV-19 via the intranasal route are protected against lower respiratory tract infection with B.1.351

Intranasal vaccination of hamsters with ChAdOx1 nCoV-19 resulted in a reduction of shedding of SARS-CoV D614G^4^. We hypothesized that a similar reduction in shedding would be found upon inoculation with B.1.351. Animals were vaccinated with 2.5 × 10^8^ IU ChAdOx1 nCoV-19/animal, either via the intranasal route or via the intramuscular route. A new stock of B.1.351 was obtained, and next gen sequencing revealed no SNPs in the spike protein (isolate hCoV-19/USA/MD-HP01542/2021) from here on referred to as B.1.351-2. Sixty days post vaccination, animals were challenged with 10^4^ TCID_50_ of B.1.351-2. As a control group, naïve animals were inoculated. Weight loss was minimal in control animals, and absent in vaccinated animals (Figure 6a). At 5 DPI, four animals per group were euthanized. Compared to control animals, lung:BW ratio of vaccinated animals was significantly reduced, and there was no difference between the two vaccine groups (Figure 6b). No viral sgRNA or infectious virus was detected in lung tissue of vaccinated animals, whereas it was abundantly present in lung tissue of control animals (Figure 6c-d). As shown in the previous experiment, no difference in the amount of virus shed was found when animals were vaccinated via the IM route. In contrast, a significant reduction was found in animals vaccinated via the IN route (Figure 6g).

**Figure 6.**
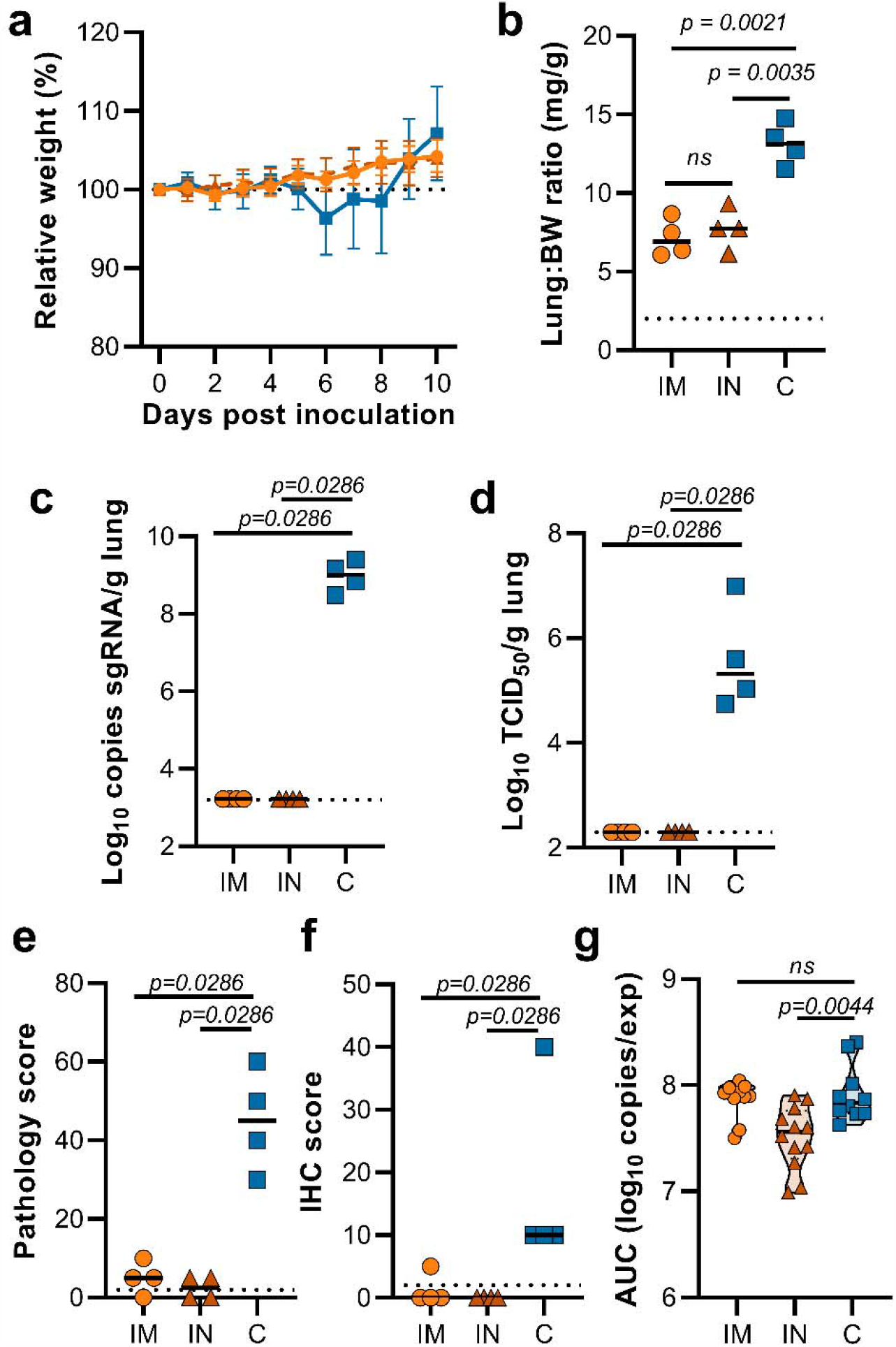
Vaccination of Syrian hamsters with ChAdOx1 nCoV-19 via the IM or IN route prevents lower respiratory tract infection with SARS-CoV-2 VOC B.1.351-2. Hamsters were vaccinated by the IM or the IN routs 60 days prior to challenge with B.1.351-2. a. Relative weight upon intranasal challenge with 10^4^ TCID_50_ of B.1.351. Shown is geometric mean with 95% confidence interval (CI). * = p-value<0.005, corrected for multiple comparisons using the Holm-Šidák correction. b. Lung:body weight (BW) ratio (mg:g) of hamsters euthanized at 5 DPI. Line = median.c. sgRNA viral load in lung tissue obtained at 5 DPI. Line = median. Dotted line = limit of detection. d. Infectious SARS-CoV-2 titer in lung tissue obtained at 5 DPI. Line = median. Dotted line = limit of detection. e. Percentage affected lung tissue per animal as determined via histology. Line = median. f. Percentage of lung tissue positive for SARS-CoV-2 antigen per animal. Line = median. g. Truncated violin plot of area under the curve (AUC) analysis of shedding as measured by sgRNA analysis in swabs collected on 1 – 5 DPI. Dashed line = median. Dotted line = quartiles. Statistical significance determined via mixed-effect analyses (a), or Mann-Whitney test (b-g). IM = ChAdOx1 nCoV-19 vaccinated via intramuscular route; IN = ChAdOx1 nCoV-19 vaccinated via intranasal route; C = naïve animals; Orange circle = Hamsters vaccinated with ChAdOx1 nCoV-19 via intramuscular route, Orange triangle = Hamsters vaccinated with ChAdOx1 nCoV-19 via intranasal route, Blue square = Naïve hamsters.

Histopathology of the lungs collected 5 DPI show pulmonary lesions of control animals with a moderate to marked broncho-interstitial pneumonia extending into the adjacent alveoli. Bronchi and bronchioles had multifocal necrotic epithelial cells and scattered to numerous infiltrating neutrophils and macrophages. Alveolar septa were expanded by edema fluid and leucocytes(Figure 7c&f). IN vaccinated animals did not show any evidence of SARS-CoV-2 pathology (Figure 7a&d). IM vaccinated animals had rare foci of interstitial pneumonia (Figure 7b&e). Immunohistochemistry using a monoclonal antibody against SARS-CoV-2 demonstrated moderate to numerous viral antigen in bronchial and bronchiolar epithelium, scattered to numerous type I and II pneumocytes as well as rare pulmonary macrophages within the control animals. The IM vaccinated animals had none to rare instances of type I and II pneumocytes and pulmonary macrophages while the IN vaccinated animals showed no evidence of viral antigen in the lungs (Figure 7g-i).

**Figure 7.**
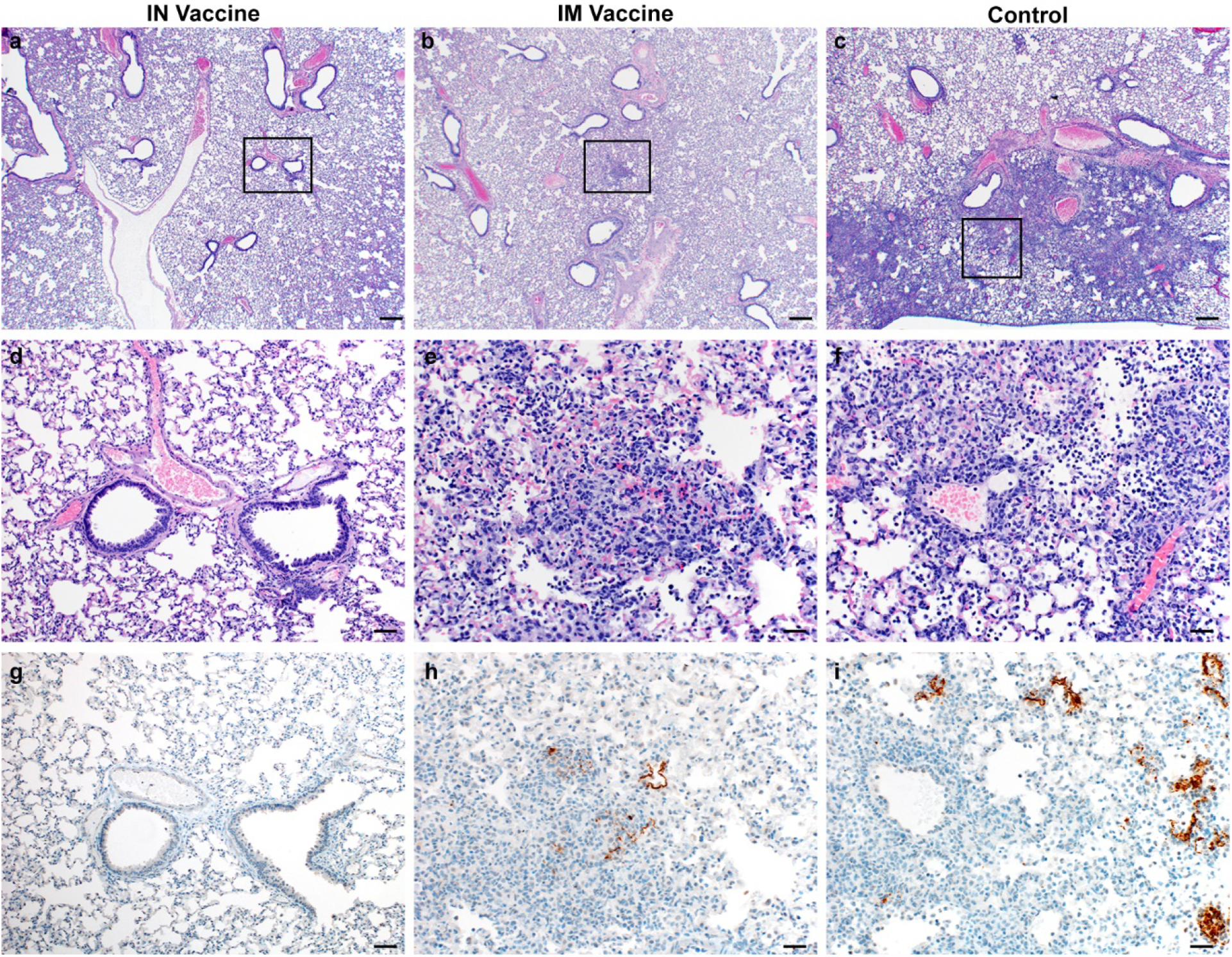
Pulmonary effects of direct IN challenge with SARS-CoV-2 variant B.1.351 in Syrian hamsters which received an IN or IM vaccine at 5 DPI. a-c: H&E 20x; a. No pathology. b-c. general prevalence of interstitial pneumonia. d-f H&E 200x; d. No pathology e. rare focus of interstitial pneumonia. f. moderate interstitial pneumonia. g-i. IHC staining against N protein SARS-CoV-2, 200x. g. No viral antigen. h. Rare foci of viral antigen in type I pneumocytes. i. Viral antigen within the larger area of interstitial pneumonia in type I pneumocytes (20x bar = 200µm; 200x bar = 20µm).

This study demonstrates efficacy of the ChAdOx1 nCoV-19 vaccine against circulating variants of concern in the SARS-CoV-2 Syrian hamster model. The Syrian hamster SARS-CoV-2 infection model is characterized by natural susceptibility to SARS-CoV-2 and development of a robust upper and lower respiratory tract infection^6^. The hamster model has been successfully used for the preclinical development of several vaccines including the Ad26 and mRNA-1273 vaccines by Janssen^7^ and Moderna^8^, respectively. Several groups have reported the effect of spike protein substitutions observed in B.1.1.7 and B.1.351 VOCs on the virus neutralizing capacity of serum obtained from vaccinated or convalescent individuals. In general, these studies conclude that the substitutions found in the B.1.1.7 spike protein have limited to no effect on virus neutralization titres^9–15^. Data from a UK phase III trial taken from a time when B.1.1.7 predominated, showed minimal impact on ChAdOx1 nCoV-19 vaccine efficacy^15^. Likewise, in an observational study of vaccine effectiveness in adults aged over 70 years in the UK, a single dose of either ChAdOx1 nCoV-19 or the Pfizer/BioNTech vaccine BNT162b2 reduced hospitalization in elderly adults with co-morbidities by 80%^16^. In contrast, the substitutions found in the B.1.351 spike protein (Table 1) result in a significant reduction of virus neutralizing capacity with pseudotype or infectious virus neutralization assays^9–15,17,18^. The ChAdOx1 nCoV-19 vaccine showed a 9 times reduction in neutralizing antibody titer against B.1.351 than against an earlier variant circulating in South Africa^10,19^. In a phase II study of ChAdOx1 nCoV-19 in South Africa, in 2000 adults with a median age of 31 years, vaccine efficacy against mild to moderate disease was reduced when the virus recovered after infection was B.1.351 (19 cases in the vaccinated group and 20 in the placebo group)^19^. Vaccine efficacy against severe disease could not be determined as no severe cases occurred in this young cohort. The South African arm of the ENSEMBLE study which tested vaccine efficacy after a single dose of Janssen’s COVID-19 vaccine candidate enrolled 6,576 participants in South Africa, out of a total of 43,783 in multiple countries, with 34% of participants across the study aged over 60 years. Vaccine efficacy against moderate to severe disease was 64% (CI 41.2%, 78.7%) in South Africa compared to 72% (CI 58.2%, 81.7%) in the USA at 28 days post vaccination^20^. Efficacy of the vaccine against severe to critical disease was 81.7% in South Africa, which was similar to the reported 85.9% and 87.6% in the USA and Brazil, respectively^20^. Vaccine efficacy against mild disease was not reported. These clinical trial results are consistent with the findings of the preclinical study reported here; ChAdOx1 nCov-19 may be less effective at reducing upper respiratory tract infection caused by B.1.351 than by B.1.1.7, consistent with reduced efficacy against mild disease. However, complete protection against lower respiratory tract disease was observed in this challenge study, consistent with protection against severe disease. Based on our data, we hypothesize that the currently available vaccines will likely still protect against severe disease and hospitalization caused by VOC B.1.351.

Limited data on the immunological determinants of protection are available, however recent data from rhesus macaques indicate that relatively low neutralizing antibody titers are sufficient for protection against SARS-CoV-2, and that cellular immune responses may contribute to protection if antibody responses are suboptimal^21^. Induction of binding and neutralizing antibodies as well as SARS-CoV-2 cellular spike protein-specific T cell responses after vaccination have been reported^22,23^ and most SARS-CoV-2 specific T cell epitopes in both convalescent and vaccinated individuals are not affected by the AA substitutions found in the spike protein of the B.1.1.7 and B.1.351 variants^24^. Protection against severe COVID-19 disease might be mediated by T cells and therefore may not be different between the current variants.

However, as T cell-mediated protection in the lower respiratory tract can only act after the initial infection has occurred, mild, polymerase chain reaction (PCR)-positive disease may still occur in the upper respiratory tract.

It should be noted that when the B.1.351 virus stock used to challenge hamsters contained two additional non-fixed AA substitutions; Q677H and R682W at 88% and 89%, respectively. The relative presence of these two AA substitutions was markedly reduced at 1 DPI and absent on 5 DPI. In addition, they were only present in lung tissue of one control hamster at 5 DPI, suggesting that they are rapidly selected against in the SARS-CoV-2 hamster model over the course of infection. Nonetheless, since the substitutions thought to be important in immune evasion, such as E484K^18^, are still present in the virus stock, efficient replication and lung pathology was observed in infected hamsters, we believe that conclusions can still be reached from the data presented in Figure 3. Additionally, we used a B.1.351 stock without AA mutations in the final experiments. Again, we did not find any disease in vaccinated hamsters inoculated with B.1.351, whether they received an IN or an IM vaccination.

Interestingly, in this same study a reduction in viral detection in oropharyngeal swabs could be detected in hamsters that received an IN vaccination, in contrast to hamsters that received an IM vaccination. These results are in line with previous studies that were done at a shorter time frame (25 and 28 days between vaccination and challenge)^4,25^. Our study shows that these differences last for at least 60 days post vaccination in hamsters, even with a VOC that has reduced the neutralizing ability of antibodies in sera.

Based on the current studies and healthcare priorities in real-world settings, we believe it is essential to focus on prevention of moderate to severe disease requiring hospitalization. We show that ChAdOx1 nCoV-19 vaccination resulted in complete protection against disease in hamsters. As implied by the data presented by Janssen^20^, viral vectored vaccines may provide substantial protection against lower respiratory tract infection caused by the B.1.351 variant and subsequent hospitalization and death. With the ongoing evolution of SARS-CoV-2, the readily available and cost-effective hamster model allows rapid evaluation of the protective efficacy of novel VOCs. In addition, it will allow rapid preclinical benchmarking of existing vaccines against preclinical vaccines with updated antigen designs.

## Acknowledgments

We would like to thank Mukul Ranjan, Sujatha Rashid, Kimberly Stemple, Alan Sutherland, Anita Mora, Kizzmekia Corbett, Barney Graham, Florian Krammer, Fatima Amanat, Victoria Avanzato, and the animal care takers for their assistance during the study. The following reagent was obtained through BEI Resources, NIAID, NIH: SARS-Related Coronavirus 2, Isolate hCoV-19/South Africa/KRISP-K005325/2020, NR-54009, contributed by Alex Sigal and Tulio de Oliveira, and SARS-Related Coronavirus 2, Isolate hCoV-19/England/204820464/2020, NR-54000, contributed by Bassam Hallis. Isolate hCoV-19/USA/MD-HP01542/2021 was obtained from Andrew Pekosz, John Hopkins Bloomberg School of Public Health.

## Author contributions

N.v.D. and V.J.M. designed the studies, S.C.G. and T.L. designed and provided the vaccine, R.J.F., N.v.D., D.R.A., C.K.Y, J.R.P., M.G.H., J.E.S., B.N.W., T.T., K.B., S.L.A., S.R, B.J.S., D.L., C.M., G.S, E.d.W., and V.J.M. performed the experiments, R.J.F., N.v.D., D.R.A., C.K.Y, J.R.P., M.G.H., J.E.S., S.L.A., C.M., G.S, and V.J.M. analyzed results, R.J.F., N.v.D and D.R.A. wrote the manuscript, all co-authors reviewed the manuscript.;

## Materials and Methods

### Ethics Statement

All animal experiments were conducted in an AAALAC International-accredited facility and were approved by the Rocky Mountain Laboratories Institutional Care and Use Committee following the guidelines put forth in the Guide for the Care and Use of Laboratory Animals 8^th^ edition, the Animal Welfare Act, United States Department of Agriculture and the United States Public Health Service Policy on the Humane Care and Use of Laboratory Animals. The Institutional Biosafety Committee (IBC) approved work with infectious SARS-CoV-2 virus strains under BSL3 conditions. Virus inactivation of all samples was performed according to IBC-approved standard operating procedures for the removal of specimens from high containment areas.

### Cells and virus

SARS-CoV-2 variant B.1.351-1 (hCoV-19/South African/KRISP-K005325/2020, EPI_ISL_678615) was obtained from Dr. Tulio de Oliveira and Dr. Alex Sigal at the Nelson R Mandela School of Medicine, UKZN. SARS-CoV-2 variant B.1.1.7 (hCoV-19/England/204820464/2020, EPI_ISL_683466) was obtained from Public Health England via BEI. SARS-CoV-2 variant B.1.351-2 (USA/MD-HP01542/2021, EPI_ISL_890360) was obtained from Andrew Pekosz at John Hopkins Bloomberg School of Public Health. Virus propagation was performed in VeroE6 cells in DMEM supplemented with 2% fetal bovine serum, 1 mM L-glutamine, 50 U/ml penicillin and 50 μg/ml streptomycin (DMEM2). VeroE6 cells were maintained in DMEM supplemented with 10% fetal bovine serum, 1 mM L-glutamine, 50 U/ml penicillin and 50 μg/ml streptomycin. Mycoplasma testing is performed at regular intervals and no mycoplasma was detected.

### Animal Experiments

ChAdOx1 nCoV-19 was formulated as previously described^26^. Four groups of 10, 4-6-week-old female Syrian hamsters (Envigo Indianapolis, IN) were vaccinated with 2.5 × 10^8^ infectious units of ChAdOx1 nCoV-19 vaccine or ChAdOx1-GFP delivered intramuscularly in two 50 µL doses into the posterior thighs 30 days prior to challenge. Five days prior to challenge a blood sample was collected via the retro-orbital plexus under isoflurane anesthesia and spun at 2000 g for 10 min to obtain serum. Two groups (10 ChAdOx1 nCoV-19 vaccinated and 10 ChAdOx1 GFP vaccinated hamsters) were challenged with 10^4^ TCID_50_/mL B.1.1.7 diluted in sterile Dulbecco’s Modified Eagle’s media (DMEM), in a 40 µL bolus delivered intranasally, one-half into each nostril. Two other groups (10 ChAdOx1 nCoV-19 vaccinated and 10 ChAdOx1 GFP vaccinated hamsters) were similarly challenged with B.1.351-1 also diluted in sterile DMEM. Weights were recorded daily until 14 DPI. Oropharyngeal swabs were collected daily in 1 mL of DMEM2 up until 5 DPI. On 5 DPI 4 animals from each group were euthanized. The lungs were excised, weighed, and photographed, and samples taken for qRT-PCR analysis, virus titrations and histopathology. The remaining six animals in each group were monitored daily for signs of disease and weighed until 14 DPI.

Two groups of 12 female Syrian hamsters (Envigo Indianapolis, IN) were vaccinated with 2.5 × 10^8^ infectious units of ChAdOx1 nCoV-19 vaccine or ChAdOx1-GFP delivered intramuscularly (IM group) in two 50 µL doses into the posterior thighs or delivered intranasaly (IN group) in 1 40 µL dose delivered equally split between each nostril 60 days prior to challenge. One group of 10 naïve female hamsters was used as a control. All three groups were challenged with 10^4^ TCID_50_/mL B.1.351-2 diluted in sterile Dulbecco’s Modified Eagle’s media (DMEM), in a 40 µL bolus delivered intranasally, equally split between each nostril. Weights were recorded daily until 14 DPI. Oropharyngeal swabs were collected daily in 1 mL of DMEM2 up until 5 DPI. On 5 DPI 4 animals from each group were euthanized. The lungs were excised, weighed, and samples taken for qRT-PCR analysis, virus titrations and histopathology. The remaining animals in each group were monitored daily for signs of disease and weighed until 14 DPI.

### Virus titration

Lung sections were weighed and homogenized in 1 mL of DMEM. Virus titrations were performed by end-point titration of 10-fold dilutions of virus swab media or tissue homogenates on VeroE6 cells in 96-well plates. When titrating tissue homogenate, the top 2 rows of cells were washed 2 times with PBS prior to the addition of a final 100 µl of DMEM2. Cells were incubated at 37°C and 5% CO2. Cytopathic effect was read 6 days later.

### Virus neutralization

Sera were heat-inactivated (30 min, 56 °C). After an initial 1:10 dilution of the sera, two-fold serial dilutions were prepared in DMEM2. 100 TCID_50_ of SARS-CoV-2 variant B.1.1.7 or B.1.351 was added to the diluted sera. After a 1hr incubation at 37°C and 5% CO_2_, the virus-serum mixture was added to VeroE6 cells. The cells were incubated for 6 days at 37°C and 5% CO_2_ at which time they were evaluated for CPE. The virus neutralization titer was expressed as the reciprocal value of the highest dilution of the serum that still inhibited virus replication. *RNA extraction and quantitative reverse-transcription polymerase chain reaction*

RNA was extracted from oropharyngeal swabs using the QiaAmp Viral RNA kit (Qiagen) according to the manufacturer’s instructions and following high containment laboratory protocols. Lung samples were homogenized and extracted using the RNeasy kit (Qiagen) according to the manufacturer’s instructions and following high containment laboratory protocols. A viral sgRNA^27^ specific assay was used for the detection of viral RNA. Five μL of extracted RNA was tested with the Quantstudio 3 system (Thermofisher) according to instructions from the manufacturer. A standard curve was generated during each run using SARS-CoV-2 standards containing a known number of genome copies.

### Viral RNA sequencing

For sequencing from viral stocks, sequencing libraries were prepared using Stranded Total RNA Prep Ligation with Ribo-Zero Plus kit per manufacturer’s protocol (Illumina) and sequenced on an Illumina MiSeq at 2 × 150 base pair reads. For sequencing from swab and lung tissue, total RNA was depleted of ribosomal RNA using the Ribo-Zero Gold rRNA Removal kit (Illumina).

Sequencing libraries were constructed using the KAPA RNA HyperPrep kit following manufacturer’s protocol (Roche Sequencing Solutions). To enrich for SARS-CoV-2 sequence, libraries were hybridized to myBaits Expert Virus biotinylated oligonucleotide baits following the manufacturer’s manual, version 4.01 (Arbor Biosciences, Ann Arbor, MI). Enriched libraries were sequenced on the Illumina MiSeq instrument as paired-end 2 × 151 base pair reads. Raw fastq reads were trimmed of Illumina adapter sequences using cutadapt version 1.12^28^ and then trimmed and filtered for quality using the FASTX-Toolkit (Hannon Lab, CSHL). Remaining reads were mapped to the SARS-CoV-2 2019-nCoV/USA-WA1/2020 genome (MN985325.1) or hCoV-19/England/204820464/2020 (EPI_ISL_683466) or hCoV-19/SouthAfrica/KRISP-K005325/2020 (EPI_ISL_678615) using Bowtie2 version 2.2.9^29^ with parameters --local --no-mixed -X 1500. PCR duplicates were removed using picard MarkDuplicates (Broad Institute) and variants were called using GATK HaplotypeCaller version 4.1.2.0^30^ with parameter –ploidy 2. Variants were filtered for QUAL > 500 and DP > 20 using bcftools.

### Expression and purification of SARS-CoV-2 S and receptor binding domain

Protein production was performed as described previously^31,32^. Expression plasmids encoding the codon optimized SARS-CoV-2 full length S and RBD were obtained from Kizzmekia Corbett and Barney Graham (Vaccine Research Center, Bethesda, USA)^33^ and Florian Krammer (Icahn School of Medicine at Mt. Sinai, New York, USA)^34^. Expression was performed in Freestyle 293-F cells (Thermofisher), maintained in Freestyle 293 Expression Medium (Gibco) at 37°C and 8% CO2 shaking at 130 rpm. Cultures totaling 500 mL were transfected with PEI at a density of one million cells per mL. Supernatant was harvested 7 days post transfection, clarified by centrifugation and filtered through a 0.22 µM membrane. The protein was purified using Ni-NTA immobilized metal-affinity chromatography (IMAC) using Ni Sepharose 6 Fast Flow Resin (GE Lifesciences) or NiNTA Agarose (QIAGEN) and gravity flow. After elution the protein was buffer exchanged into 10 mM Tris pH8, 150 mM NaCl buffer (S) or PBS (RBD) and stored at − 80°C.

### ELISA

ELISA was performed as described previously^26^. Briefly, maxisorp plates (Nunc) were coated overnight at 4°C with 50 ng/well S or RBD protein in PBS. Plates were blocked with 100 µl of casein in PBS (Thermo Fisher) for 1hr at RT. Serum diluted 1:6,400 was further 2-fold serially diluted in casein in PBS was incubated at RT for 1hr. Antibodies were detected using affinity-purified polyclonal antibody peroxidase-labeled goat-anti-monkey IgG (Seracare, 074-11-021) in casein followed by TMB 2-component peroxidase substrate (Seracare, 5120-0047). The reaction was stopped using stop solution (Seracare, 5150-0021) and read at 450 nm. All wells were washed 4x with PBST 0.1% tween in between steps. Threshold for positivity was set at 2x OD value of negative control (serum obtained from unvaccinated hamsters prior to start of the experiment).

### Data availability statement

Data have been deposited in Figshare 10.6084/m9.figshare.14210879.

